# Construction of Accurate Turtle Cerebellum Models and their use in Simulating Neuromodulation using the BEM-FMM

**DOI:** 10.1101/2025.04.08.647756

**Authors:** Gregory M. Noetscher, Sam Small-Zlochower, Padmavathi Sundaram, Sergey N. Makaroff

## Abstract

Digital representations of Purkinje cells from turtle cerebellum, which have long been to study cellular transmission dynamics and neuroplasticity, are necessary to conduct simulations of neuromodulation methods, including Transcranial Magnetic Stimulation (TMS). *Methods:* Data collected using microCT with 0.6 um isometric resolution are processed, segmented, and arranged to enable high resolution simulation via the Boundary Element Fast Multipole Method. *Results:* Fully computer aided design compatible, manifold triangular surface meshes were generated from microCT data and used to simulate cellular electromagnetic response to TMS. *Conclusion:* Accurate Purkinje cells may be readily generated and manipulated to create representative models of turtle cerebellum. *Significance:* The techniques employed herein may be used to generate accurate surface mesh of Purkinje cells which can enable computational electromagnetics code verification and serve as the basis for skeletal models used for bidomain biophysical simulations in tools such as NEURON to study neurostimulation dynamics and neuromodulation.

## I. Introduction

PURKINJE cells in the cerebellum have been thoroughly studied since the mid-1960s due to their relatively simple structure, neuroplasticity, and synaptic physiology [1]-[3]. Turtle Purkinje cells are particularly well suited for long duration *in-vitro* experiments due to their robust electrophysiological properties and ability to survive under hypoxic conditions [4]-[13]. The combination of these attributes has enabled the use of turtle cerebellum to experimentally characterize the response at the cellular level to invasive and non-invasive neuromodulation techniques, including Transcranial Magnetic Stimulation (TMS) [14], allowing for the validation of relevant model construction and related simulations.

This paper provides the methods used for creation of anatomically realistic, computer aided design (CAD) compatible models of turtle Purkinje cells, including microCT (uCT) data collection, image processing, and triangular surface mesh creation. Once generated, these models were used to simulate via the Boundary Element Fast Multipole Method (BEM-FMM) [15]-[17] their response to an external field applied via TMS.

## II. Methods

### A. Data collection

Intact cerebellum of turtle (Pseudemys scripta elegans) was extracted using a surgical procedure approved by the Institutional Animal Care and Use Committee (IACUC). The tissue was fixed and stained via the Golgi-Cox impregnation method using the FD Rapid GolgiStain Kit (FD NeuroTechnologies, Inc. MD) [18] and then prepared for imaging using micro-computed tomography (microCT). The samples were imaged using microCT at a resolution of 0.6 um (isotropic). The sample was thereby characterized by 961 images, each consisting of 906×993 pixels. It should be noted that Purkinje cell axons are not obtained using this process.

### B. Image Processing

The resulting image stack was imported into Fiji [19], an enhancement of the popular ImageJ utility [20], which hosts sophisticated suite of tools for further processing to facilitate segmentation. As can be seen in the top section of Fig. 1, pixels with the highest intensity represent Purkinje cell structures including the soma and dendrites. The surrounding ‘background’ pixels represent noise in the sample and should be eliminated to maximize the contrast between desirable and undesirable features for segmentation.

**Fig. 1.**
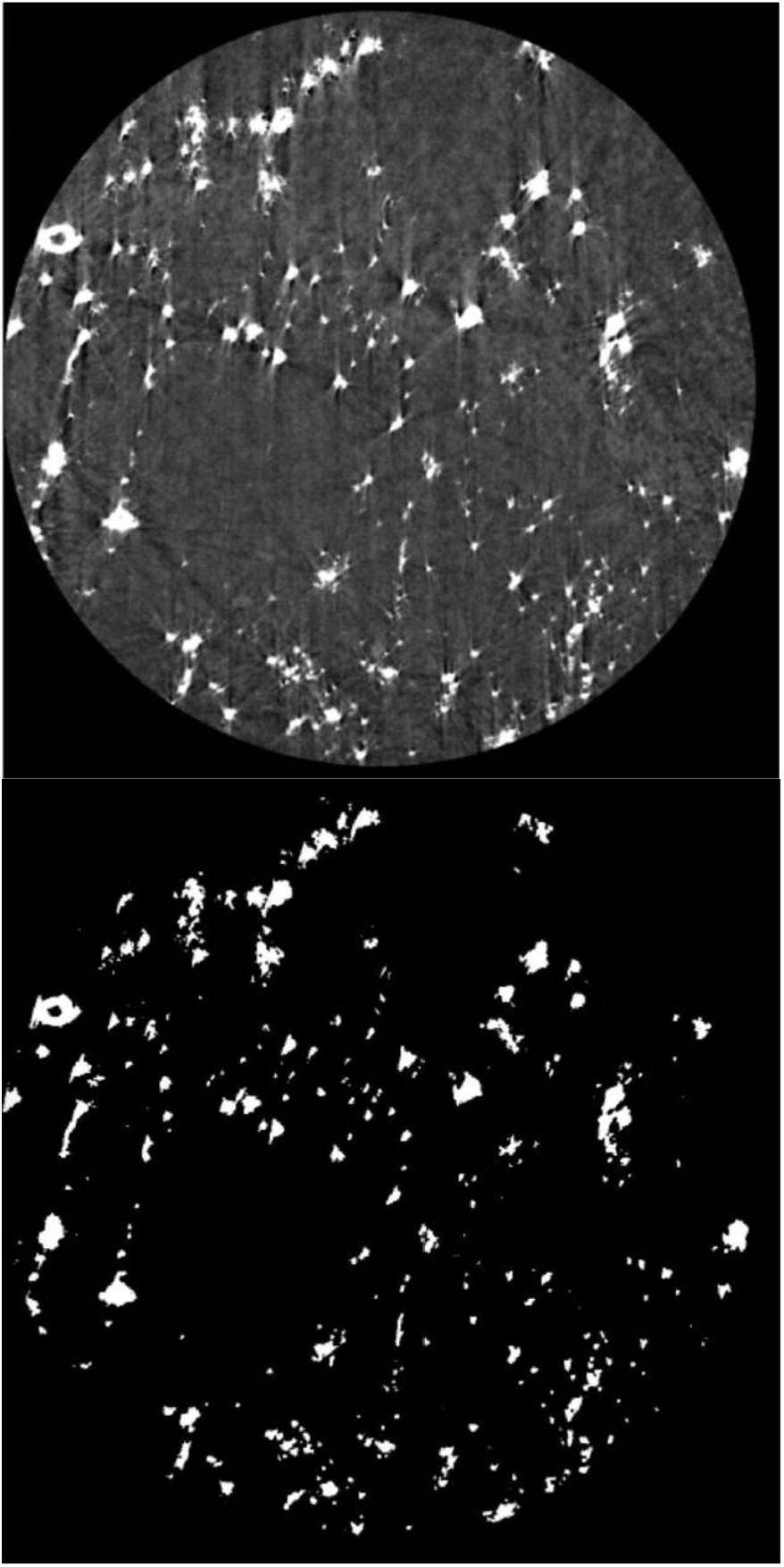
Top – Sample image of microCT data generated with an isotropic resolution of 0.6 µm; Bottom – Processed image prepared for segmentation. Note the areas of high contrast representing cell structures.

The average pixel intensity value of the sample background was calculated and subtracted from the image stack. While this improved the contrast, the result contained a significant amount of noise that complicated the segmentation process.

Fiji’s ‘Auto Threshold’ tool was investigated to ascertain its performance in developing a binarized version of the original dataset. Algorithms in the Auto Threshold tool include among others fuzzy thresholding, bimodal histogram smoothing, and minimum and maximum entropy thresholding. Of those tested, the Intermodes algorithm [21] provided the best balance between removing the surrounding background noise and preserving the cell structures of interest. The results of this processing can be seen at the bottom of Figure 1 and were used for cell segmentation.

### C. Segmentation

Image segmentation was carried out using ITK-Snap, an open-source platform optimized for medical data that provides simultaneous viewing of segmented structures in three orthogonal planes [22]. Given the binary nature of the processed image stack, semi-automatic segmentation using an active contour method within ITK-SNAP was suitable for generating full segmentations of individual Purkinje cells, as shown in Figure 2. ITK-SNAP can directly export an individual segmentation as a triangular surface mesh file in the STL format, which is ideally suited BEM-FMM simulation.

**Fig. 2.**
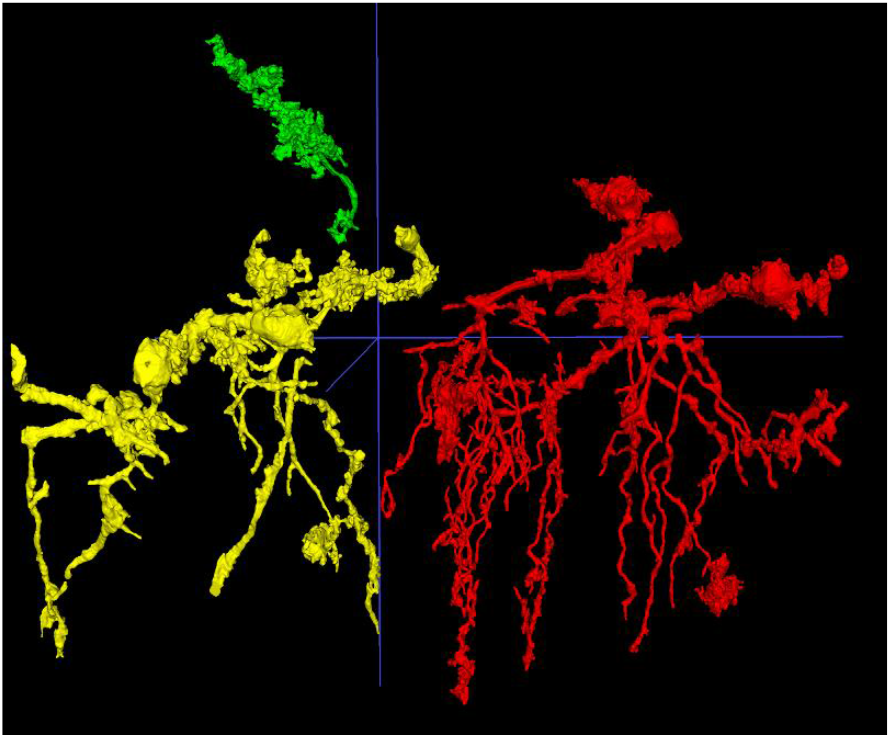
Example cell segmentations obtained from ITK-SNAP. Each color represents a single Purkinje cell, saved as an individual mesh file.

### D. Simulation

Simulations were conducted using a BEM-FMM based computational electromagnetic platform built in MATLAB [17]. The underlying methodology is a charge-based formulation of the Boundary Element Method – charges induced as a result of a primary stimulus field are calculated on model surfaces. These induced charges tend to alter the primary stimulus field, producing the total field anywhere in space, as governed by Coulomb’s law. The Fast Multipole Method is employed to accelerate the calculation of electric potential integrals. This formulation results in very high accuracy and essentially unconstrained numerical field resolution and is ideally suited for calculations on complex, entangled structures.

## III. Results and Discussion

The image processing and segmentation pipeline previously introduced produced Purkinje cells from the turtle cerebellum with dimensions consistent with expected values. Single cells have been simulated to examine the charge accumulation due to the application of an external electric field via TMS. The diameters of individual cell somas were approximately 30 µm and revealed extensive dendritic arborization with diameters of about 5 µm. For uniformity, cell somas were removed and replaced with spheres.

Due to the preparation process used on the sample, not all Purkinje cells are visible for segmentation. Therefore, a number of representative cells were duplicated to generate a model with the appropriate cell density of 100 Purkinje cells per millimeter of cerebellar cortex length. These cells were arranged in a single layer and given random orientations. An example of this construction is shown in Figure 3.

**Fig. 3.**
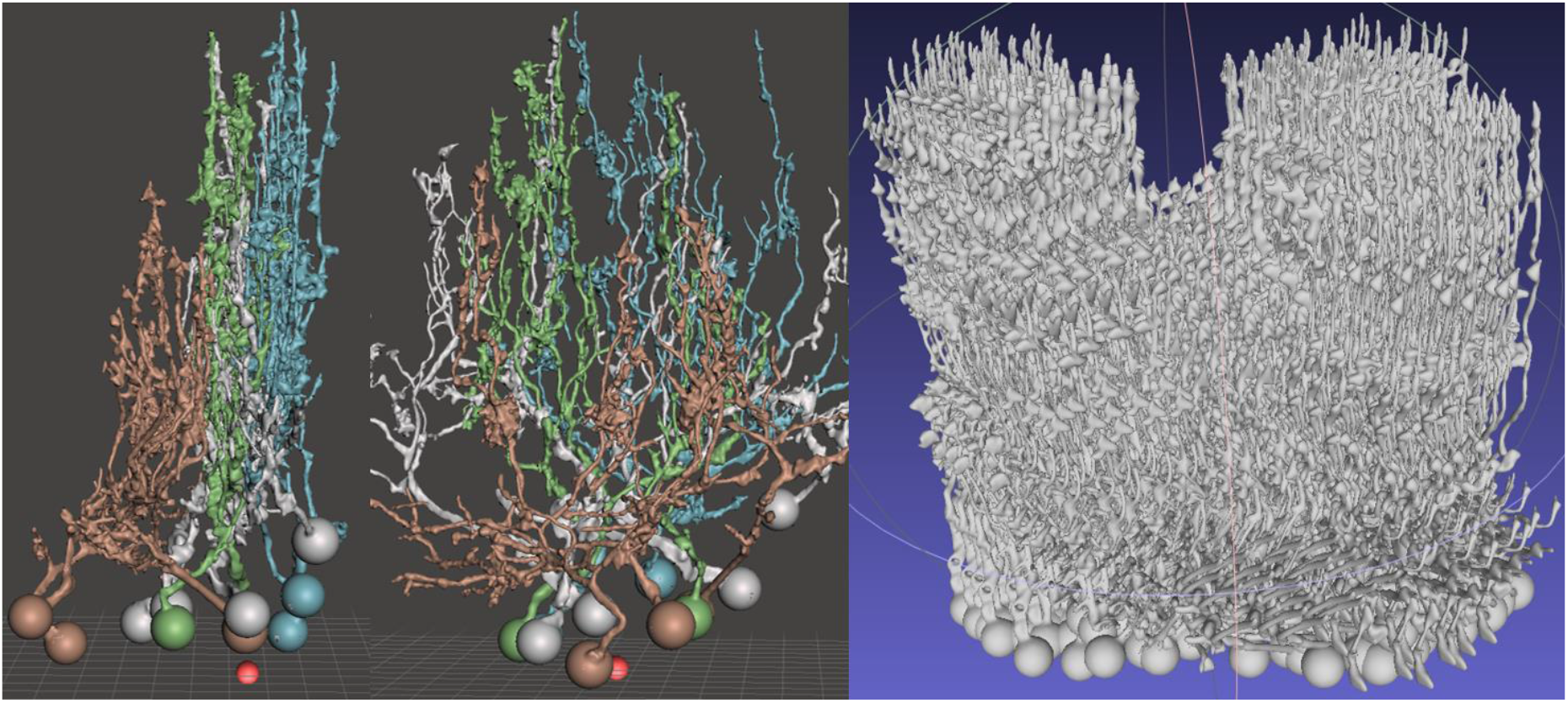
Representative groupings employed to generate appropriate cerebellar cell cortex density.

When viewed as a collection, the accumulated charge on cell surfaces has a distorting impact on the impressed electric field. This impact may modify the activating threshold of an individual cell and provide guidance on the design of appropriate neuromodulation protocols.

## IV. Conclusion

A workflow designed to enable the extraction of Purkinje cells from turtle cerebellum data has been demonstrated, resulting in accurate reconstruction of models characterizing the physical form of the cell. These models have been integrated into a high-resolution BEM-FMM based simulation platform enabling fast modeling of TMS induced charge accumulation and corresponding electric field distributions on cells.

## Acknowledgment

The authors wish to thank Dr. Chunling Dong from the Martinos Center for Biomedical Imaging, Massachusetts General Hospital, and Harvard Medical School, Boston and Dr. Richard Schalek from the Lichtman Lab at Harvard University for preparation and imaging of the turtle cerebellum samples.

